# SLFN-generated 3′-truncated tRNA^Leu^ or tRNA^Ser^ together with tRNase Z^L^ works as a sequence-specific RNA cutter

**DOI:** 10.64898/2026.04.12.718003

**Authors:** Masayuki Takahashi, Masayuki Nashimoto

## Abstract

A complex of the 3′-truncated tRNA^Arg^ that lacks 9 nt and tRNase Z^L^ works as a GCCC-recognizing RNA cutter. It recognizes an RNA substrate via four Watson-Crick-Franklin base-pairings with the 3′-truncated tRNA^Arg^. Human SLFN11 and SLFN13 can generate 3′-truncated tRNA^Leu^ that lacks 10 nt and 3′-truncated tRNA^Ser^ that lacks 11 nt, respectively, from their corresponding mature tRNAs. Here, we investigated if these 3′-truncated tRNAs together with tRNase Z^L^ work as sequence-specific RNA cutters. We examined five RNA targets for cleavage by recombinant human tRNase Z^L^ in the presence of the 3′-truncated tRNA^Leu^ or tRNA^Ser^. We demonstrated that the 3′-truncated tRNA^Leu^ and tRNA^Ser^ together with tRNase Z^L^ indeed work as ∼6-base-recognizing and 7-base-recognizing RNA cutters, respectively.

## Introduction

In 1989, we have happened to find a 4-nucleotide (nt) recognizing RNA cutter in mouse cell extracts and have named it RNase 65 [1–4]. RNase 65 has turned out to be a complex of tRNA 3′-processing endoribonuclease (tRNase Z^L^) and 3′-truncated tRNA that lacks 9 nt, and it recognizes an RNA substrate via four Watson-Crick-Franklin base-pairings with the 3′-truncated tRNA [5]. Specifically, the 5′-terminal sequence 5′-GGGC-3′ of 3′-truncated tRNA^Arg^ base-pairs with the sequence 5′-GCCC-3′ of substrate RNA, while the 5′-terminal sequence 5′-GGGG-3′ of 3′-truncated tRNA^Ala^ base-pairs with the sequence 5′-CCCC-3′ of substrate RNA. We have also detected 3′-truncated tRNA^Gln^, tRNA^Gly^ and tRNA^Met^ in the mouse cell extracts [2], and 3′-truncated tRNA^Gly^ and tRNA^Val^ have been found in mouse epididymis [6]. The physiological role and the genuine cellular substrate RNA of RNase 65 remain to be elucidated. How 3′-truncated tRNA can be generated also remains unsolved. However, since 3′-truncated tRNA^Arg^can be generated by inducing apoptosis in human cells [7], we suppose that the RNase 65 activity may be involved in RNA catabolism.

While attempting to elucidate the unsolved RNase 65 issues, we also have been studying the interactions of tRNase Z^L^ with various RNA substrates and developing a gene suppression method by modifying the RNase 65 activity [8]. In the course of these extensive studies, in 2007, we also have happened to find 5′-half-tRNA^Glu^ in HEK293 cells as a tRNase Z^L^-bound RNA, and have shown that 5′-half-tRNA^Glu^ downregulates the PPM1F mRNA in the cells [7]. After this discovery, the existence of many other 5′-half-tRNAs and many different-type tRNA fragments in mammalian cells has been uncovered, and various modes of action of these tRNA fragments in gene regulation are suggested [9,10].

Recently, we have learned that human SLFN11 and SLFN13 among six members of the SLFN gene family on chromosome 17 can generate 3′-truncated tRNA^Leu^ that lacks 10 nt and 3′-truncated tRNA^Ser^ that lacks 11 nt, respectively, from their corresponding mature tRNAs [11,12]. These observations naturally led us to wonder if these SLFN-generated 3′-truncated tRNAs together with tRNase Z^L^ work as sequence-specific RNA cutters and if the 3′-truncated tRNA^Arg^ that lacks 9 nt is generated by one of the SLFN gene family members. In this study, we addressed the former question using *in vitro* synthesized 3′-truncated tRNAs and recombinant human tRNase Z^L^, and demonstrated that the 3′-truncated tRNA^Leu^ or the 3′-truncated tRNA^Ser^ together with tRNase Z^L^ shows an endoribonuclease activity with their own sequence specificity.

## Materials and methods

### *In vitro* 3′-truncated tRNA synthesis

3′-truncated tRNA^Leu^ (5′-GCCAGGAUGGCCGAGUGGUUAAGGCGUUGGACUUAAGAUCCAAUGGACAUAUGUCCGCGUGGGUUCGAACCCCACU-3′) [11] and 3′-truncated tRNA^Ser^ (5′-GUAGUCGUGGCCGAGUGGUUAAGGCGAUGGACUUGAAAUCCAUUGGGGUUUCCCCGCGCAGGUUCGAAUCCUGC-3′) [12] were synthesized *in vitro* with T7 RNA polymerase (Takara, Shiga, Japan) from the corresponding synthetic DNA templates. The reactions were performed under the conditions recommended by the manufacturer (Takara), and the 3′-truncated tRNAs were purified by denaturing gel electrophoresis [7].

### Preparation of recombinant human tRNase Z^**L**^

The histidine-tagged human tRNase Z^L^ that lacks N-terminal 30 amino acids was overexpressed using the plasmid pQE-80L in *E. coli* strain Rosetta(DE3)pLysS and subsequently purified with nickel-agarose beads [7].

### Target RNA synthesis

Five 26-nt 5′-FAM-labeled RNA targets, Target-1 (5′-

GGUGUCCUGGCGACUACGGUGUAAGC-3′), Target-2 (5′-

GGUGAGCUGGCAACUACGGUGUAAGC-3′), Target-3 (5′-

GGUGACCUGGCAAACUACGGUGUAAG-3′), Target-4 (5′-

GGUGAGAUGGCAACCUACGGUGUAAG-3′), and Target-5 (5′-

GGUGUCCAGGCGAAUACGGUGUAAGC-3′) were chemically synthesized and subsequently purified by high-performance liquid chromatography by Nippon Bioservice (Saitama, Japan).

### RNA cleavage assay

*In vitro* tRNase Z^L^ cleavage assay was performed at 37 °C with 2 pmol of 5′-FAM-labeled target RNA, 4 pmol of 3′-truncated tRNA and 2.5 pmol of recombinant human tRNase Z^L^ in a 20-μl mixture containing 10 mM Tris–HCl (pH 7.1) and 1–15 mM MgCl_2_ or 0.1–1 mM spermidine. The reaction was started by adding tRNase Z^L^ to the reaction mixture that was pre-incubated at room temperature for 5 min. The reaction products were resolved on a 20% polyacrylamide gel containing 8 M urea and analyzed using a Gel Doc EX imager (Bio-Rad, Tokyo, Japan).

### Estimation of observed reaction rate constants

An observed reaction rate constant (*k*_obs_) for target RNA cleavage by tRNase Z^L^ in the presence of 5 mM MgCl_2_ or 0.2 mM spermidine was estimated from amounts of the cleaved product (P) after 15-, 30-, 60- and 90-min reactions. A *k*_obs_ value and an effective substrate amount (S0_eff_) were calculated by curve fitting of the experimental data to the equation P = S0_eff_ × (1 − exp (− *k*_obs_ × time)) using the scipy.optimize module in Python.

### Results

The 3′-truncated tRNA^Leu^ that lacks 10 nt and the 3′-truncated tRNA^Ser^ that lacks 11 nt were synthesized by *in vitro* T7 RNA polymerase transcription of their corresponding synthetic DNA templates. The first nucleotide A of the 3′-truncated tRNA^Leu^ was replaced by G due to demand of efficiency of T7 RNA polymerase transcription. The five 26-nt RNA targets Target-1–Target-5 were chemically synthesized with 5′-FAM labeling. Target-1 contains two 7-nt sequences that are fully complementary to those of the 5′-acceptor-stems of tRNA^Leu^ and tRNA^Ser^, while each of Target-2–Target-5 contains one to three nt mismatches with the 5′-acceptor-stem sequences. Potential secondary structures of the complexes between 3′-truncated tRNAs and RNA targets are shown in Fig. 1.

**Fig. 1.**
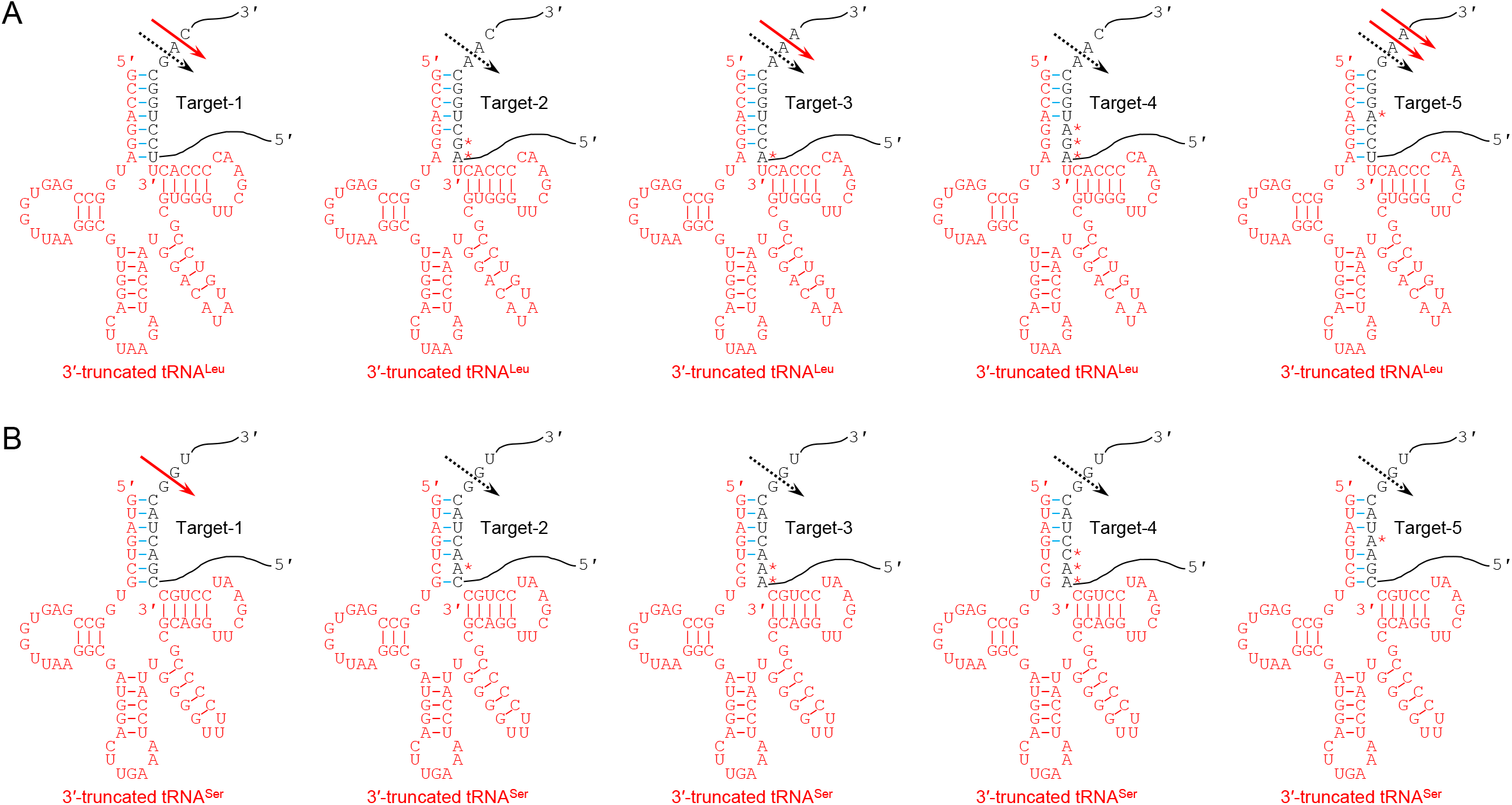
Possible secondary structures of 3′-truncated tRNA/target RNA complexes. (A) Possible secondary structures of complexes of 3′-truncated tRNA^Leu^ with Target-1–Target-5. (B) Possible secondary structures of complexes of 3′-truncated tRNA^Ser^ with Target-1–Target-5. A base in target RNA that mismatches with the 5′-acceptor-stem sequences is indicated by a red asterisk. A black dotted arrow and a red arrow denote an expected cleavage site and an actual cleavage site, respectively.

We examined the five RNA targets for cleavage by recombinant human tRNase Z^L^ in the presence of the 3′-truncated tRNA^Leu^ or the 3′-truncated tRNA^Ser^. The reactions were carried out at 37 °C for 30 min in the presence of 5 mM Mg^2+^. Target-1 was cleaved in the presence of the 3′-truncated tRNA^Leu^ and tRNA^Ser^, generating ∼10-nt and ∼20-nt products, respectively (Fig. 2A). Independently of the presence of the 3′-truncated tRNAs, an ∼10-nt product of Target-1 was also observed, which would have been generated by the property of tRNase Z^L^ that it can cleave even unstructured single-stranded RNA [13]. Target-3 and Target-5, which contain one nt mismatch with the 5′-acceptor-stem sequence of tRNA^Leu^, were also cleaved in the presence of the 3′-truncated tRNA^Leu^ generating an ∼10-nt product, but not in the presence of the 3′-truncated tRNA^Ser^ (Fig. 2A). The other targets Target-2 and Target-4 were not cleaved in the presence of either 3′-truncated tRNAs (Fig. 2A).

**Fig. 2.**
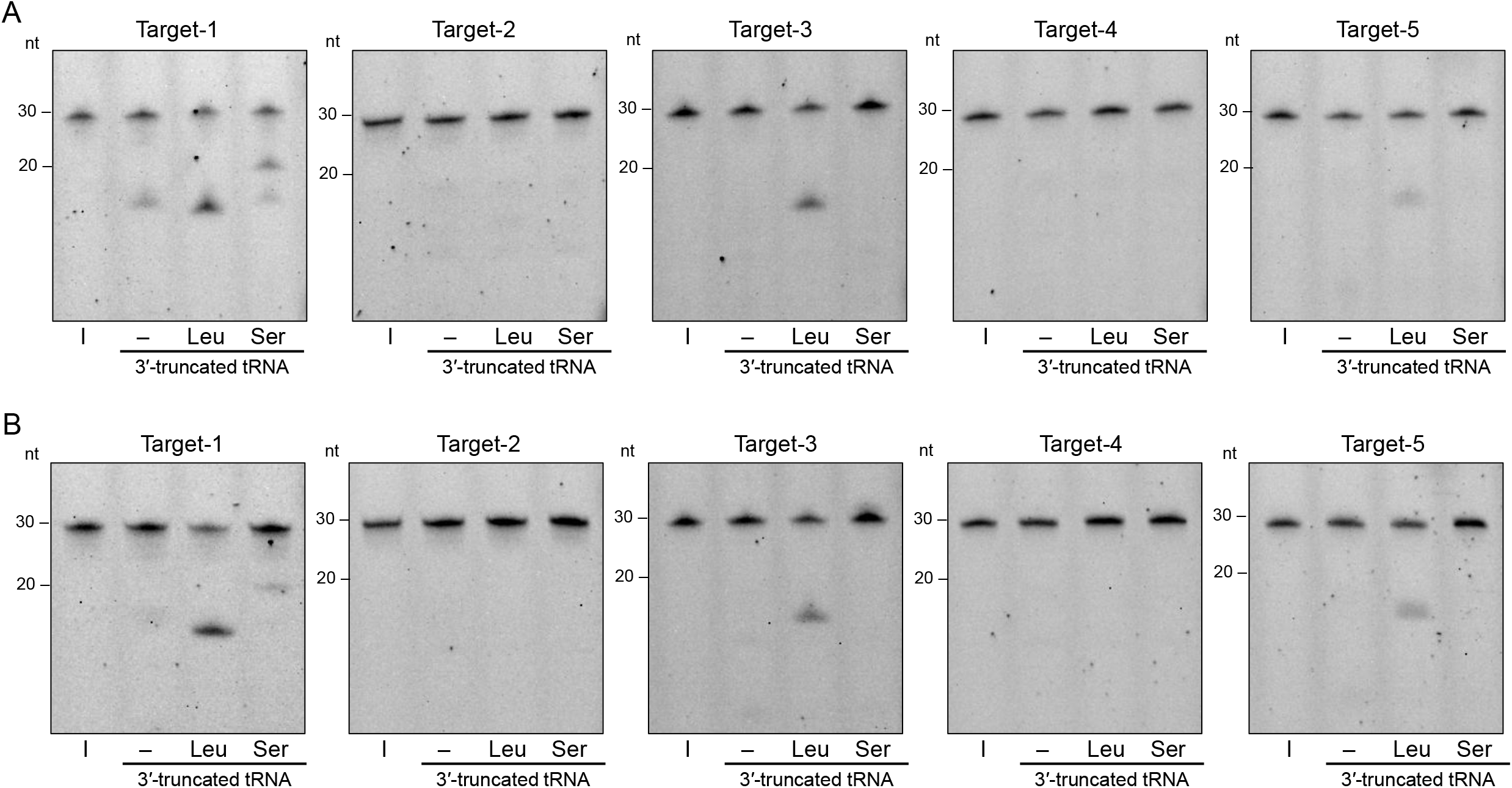
*In vitro* RNA cleavage assays with 5 mM MgCl_2_ (A) and with 0.1 mM spermidine (B). Target-1–Target-5 were incubated with recombinant human tRNase Z^L^ in the absence and presence of 3′-truncated tRNA^Leu^ and 3′-truncated tRNA^Ser^ at 37 °C for 30 min. The reaction products were analyzed by denaturing 20% polyacrylamide gel electrophoresis. Positions of the 20- and 30-nt size standards are shown on the left side of each panel. I, input RNA.

We also tested the RNA targets for cleavage by tRNase Z^L^ in the presence of mM spermidine. In the same way as the reactions with 5 mM Mg^2+^, Target-1 was cleaved in the presence of the 3′-truncated tRNA^Leu^ and tRNA^Ser^, and Target-3 and Target-5 were cleaved only in the presence of the 3′-truncated tRNA^Leu^ (Fig. 2B). Target-2 and Target-4 were not cleaved in the presence of either 3′-truncated tRNAs (Fig. 2B). Taken together, these observations suggest that the RNA cutter with the 3′-truncated tRNA^Leu^ needs at least 6 base matches with target RNA in a 7-nt consecutive sequence for the target RNA recognition and that the RNA cutter with the 3′-truncated tRNA^Ser^ needs 7 base full matches with target RNA in a 7-nt consecutive sequence (Fig. 1).

With respect to the reactions for Target-1, we determined optimal Mg^2+^ and spermidine concentrations. The reactions of Target-1 cleavage by tRNase Z^L^ were optimal at 5–10 mM Mg^2+^ both in the presence of the 3′-truncated tRNA^Leu^ and tRNA^Ser^, while they were optimal at 0.2–0.4 mM spermidine both in the presence of the 3′-truncated tRNA^Leu^ and tRNA^Ser^ (Fig. 3).

**Fig. 3.**
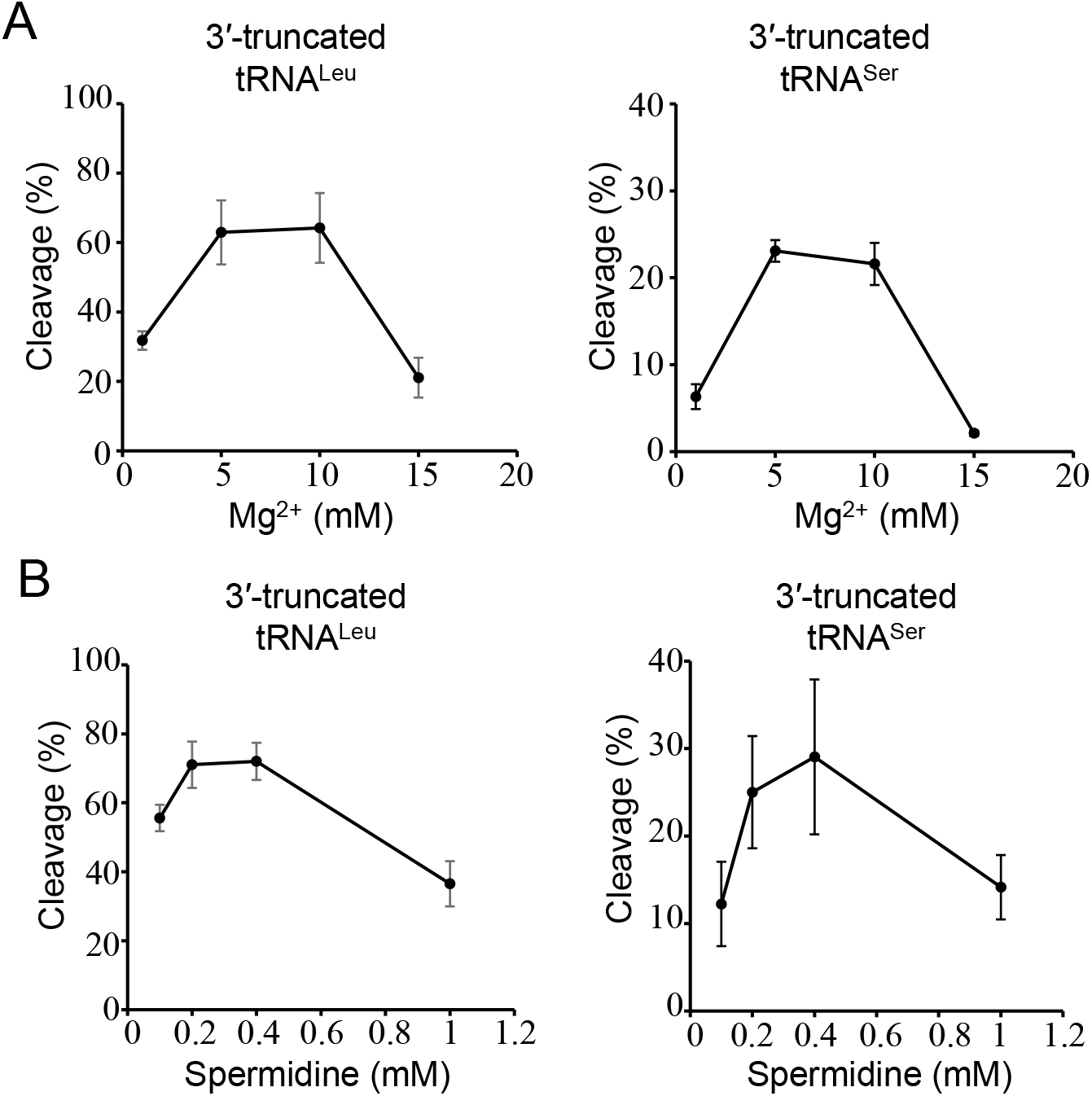
Determination of optimal Mg^2+^ and spermidine concentrations for the reactions of Target-1 cleavage by tRNase Z^L^ in the presence of the 3′-truncated tRNA^Leu^ or tRNA^Ser^. The reactions were carried out at 37 °C for 30 min with 1– 15 mM MgCl_2_ (A) or 0.1–1 mM spermidine (B). Values are mean ± SD for three replicates.

Next, we determined the exact cleavage sites of the RNA targets in the above reactions by analyzing a larger gel with an alkaline ladder as size standards. In the presence of the 3′-truncated tRNA^Leu^, Target-1 was cleaved after the 13th nt A, which is immediately downstream to the nt corresponding to a discriminator, both with 5 mM Mg^2+^ and with 0.2 mM spermidine, whereas, in the presence of the 3′-truncated tRNA^Ser^, it was cleaved after the 18th nt G, which corresponds to a discriminator, under the two conditions (Fig. 4A). In the presence of the 3′-truncated tRNA^Leu^, Target-3 was cleaved after the 13th nt A, which is immediately downstream to the nt corresponding to a discriminator, under both conditions and Target-5 was cleaved after the 14th nt A and the 13th nt A, which is immediately downstream to the nt corresponding to a discriminator, under both conditions (Fig. 4B,C).

**Fig. 4.**
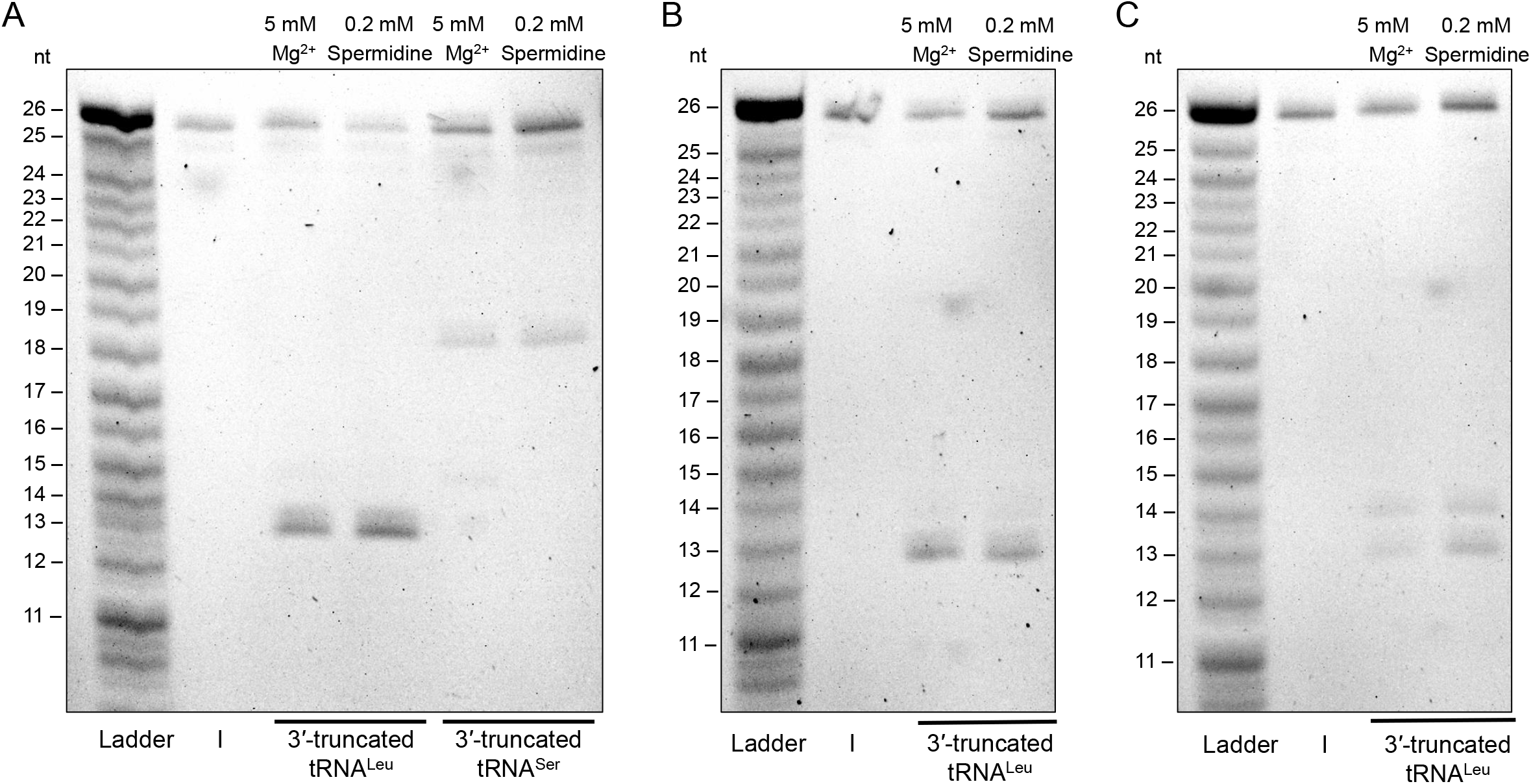
Determination of the exact cleavage sites of Target-1 (A), Target-3 (B) and Target-5 (C). The cleavage reaction was carried out in the presence of the 3′-truncated tRNA^Leu^ or tRNA^Ser^ at 37 °C for 30 min with 5 mM MgCl_2_ or 0.2 mM spermidine. The products were analyzed on a denaturing 20% polyacrylamide large gel. An alkaline ladder was prepared by incubating 5′-FAM-labeled target RNA in a solution containing 50 mM NaOH and 1 mM EDTA at 95 °C for 4 min. I, input RNA.

Lastly, with respect to the reactions of target RNA (2 pmol), 3′-truncated tRNA (4 pmol) and tRNase Z^L^ (2.5 pmol) whereby cleavage occurred, we estimated an observed reaction rate constant (*k*_obs_) and an effective substrate amount (S0_eff_). *k*_obs_ would reflect both the affinity of substrate and enzyme and the catalytic rate constant, while S0_eff_ would reflect the amount of a target RNA/3′-truncated tRNA complex that is recognizable by tRNase Z^L^.

In the cleavage reactions with Mg^2+^, the *k*_obs_ value for the reaction of Target-1 in the presence of the 3′-truncated tRNA^Leu^ was 0.047 min^−1^, whereas those for the other reactions were 0.074–0.086 min^−1^ (Fig. 5A). The S0_eff_ for the reactions of Target-1 and Target-3 in the presence of the 3′-truncated tRNA^Leu^ were high (1.5 and 0.96 pmol, respectively), while those for the reaction of Target-1 in the presence of the 3′-truncated tRNA^Ser^ and the reaction of Target-5 in the presence of the 3′-truncated tRNA^Leu^ were low (0.44 and 0.56 pmol, respectively).

**Fig. 5.**
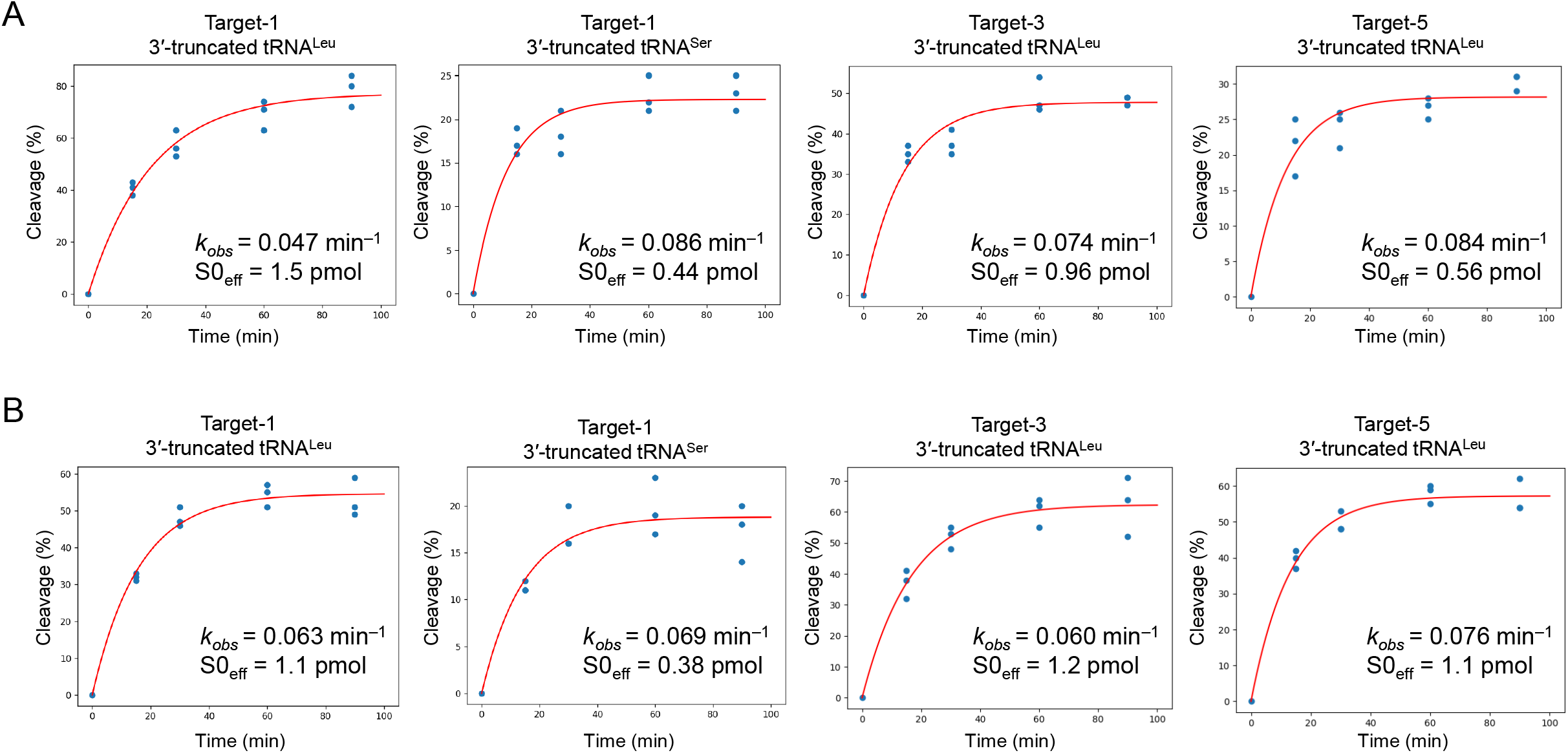
Estimation of an observed reaction rate constant (*k*_obs_) and an effective substrate amount (S0_eff_) for target RNA cleavage by tRNase Z^L^ in the presence of the 3′-truncated tRNA^Leu^ or tRNA^Ser^. The reactions were carried out at 37 °C for 15, 30, 60 and 90 min with 5 mM MgCl_2_ (A) or 0.2 mM spermidine (B).

In the reactions with spermidine, the *k*_obs_ values were 0.060–0.076 min^−1^ (Fig. 5B). The S0_eff_ for the reactions in the presence of the 3′-truncated tRNA^Leu^ were 1.1–1.2 pmol, and that for the reaction in the presence of the 3′-truncated tRNA^Ser^ was 0.38 pmol.

## Discussion

We demonstrated that the 3′-truncated tRNA^Leu^ and the 3′-truncated tRNA^Ser^ together with tRNase Z^L^ show an ∼6-base-recognizing endoribonuclease activity and a 7-base-recognizing endoribonuclease activity, respectively, under the conditions of an appropriate concentration of MgCl_2_ or spermidine. Like RNase 65, tRNase Z^L^ would also be able to form a binary complex with the 3′-truncated tRNA^Leu^ or the 3′-truncated tRNA^Ser^ [3]. It is also likely that many other 3′-truncated tRNA species that are generated by SLFNs or other nucleases together with tRNase Z^L^ can show an endoribonuclease activity with their own sequence specificity. It would be useful to name a tRNase Z^L^/3′-truncated tRNA complex RNase 65 and to add a one-letter amino-acid code according to a tRNA species such as RNase 65L and RNase 65S to specify the 3′-truncated tRNA species as proposed before [5].

It is expected that tRNase Z^L^ cleaves pre-tRNAs after their discriminator nucleotide and pre-tRNA-like complexes after a nucleotide corresponding to the discriminator nucleotide [8]. In the presence of the 3′-truncated tRNA^Ser^, Target-1 was cleaved by tRNase Z^L^ at the expected site under the two conditions, whereas, in the presence of the 3′-truncated tRNA^Leu^, Target-1 was cleaved at the site 1-nt downstream to the expected one (Fig. 1). In the presence of the 3′-truncated tRNA^Leu^, Target-3 was also cleaved at the site 1-nt downstream to the expected one, and Target-5 was cleaved both at the sites 1-nt and 2-nt downstream to the expected one. Such cleavage site shifts are often observed in the *in vitro* tRNase Z^L^ system [14,15].

Although the differences in *k*_obs_ and S0_eff_ among the reactions of Target-1, Target-3 and Target-5 with spermidine in the presence of the 3′-truncated tRNA^Leu^ were small, for the reactions with Mg^2+^, the *k*_obs_ value (0.047 min^−1^) for the reaction of Target-1 was much smaller than those of the reactions of Target-3 (0.074 min^−1^) and Target-5 (0.084 min^−1^), and S0_eff_ differed among those three reactions (Fig. 5). These observations suggest that Mg^2+^ ions affect differentially both on the formation of the 3′-truncated tRNA^Leu^/target RNA complexes and on the formation of tRNase Z^L^/substrate RNA complexes. Although there were no big differences in *k*_obs_ and S0_eff_ between the reactions of Target-1 in the presence of the 3′-truncated tRNA^Ser^ with Mg^2+^ and with spermidine, S0_eff_ for the reactions were much smaller than those for the reactions of Target-1 in the presence of the 3′-truncated tRNA^Leu^ both with Mg^2+^ and with spermidine (Fig. 5). This difference would be partly due to the difference in the number of the GC pair between 3′-truncated tRNA and target RNA (Fig. 1).

In the current study, we did not address the hard questions: What are the physiological roles of those sequence-specific RNA cutters? What are their genuine cellular target RNAs? If these RNA cutters have some physiological role, the 3′-truncated tRNA^Leu^ and tRNA^Ser^ would need to exist somewhat stably in the cells, and the conditions in the intracellular milieu would need to be appropriate for their RNA cleavage activity. If these hold, the analysis for changes in the transcriptome by introducing the 3′-truncated tRNA^Leu^ or tRNA^Ser^ into the cells would provide a clue to elucidate the physiological roles and the genuine cellular target RNAs of the sequence-specific RNA cutters.

## Acknowledgements

We thank a former lab member, Hirotaka Shonen Shibata for providing us the information on SLFN-generated 3′-truncated tRNA that motivated this study.

## Data accessibility

All the data that support the finding of this study are within the manuscript.

## Author contributions

MT: data curation, formal analysis, investigation, writing (review). MN: conceptualization, data curation, formal analysis, project administration, supervision, writing (original draft, editing).

